# Human genomic DNA is widely interspersed with i-motif structures

**DOI:** 10.1101/2022.04.14.488274

**Authors:** Cristian David Peña Martinez, Mahdi Zeraati, Romain Rouet, Ohan Mazigi, Brian Gloss, Chia-Ling Chan, Tracy M. Bryan, Nicole M. Smith, Marcel E. Dinger, Sarah Kummerfeld, Daniel Christ

## Abstract

DNA i-motif structures are formed in the nucleus of human cells and are believed to provide critical genomic regulation. While the existence of i-motif structures in human cells has been demonstrated by immunofluorescent staining and by characterisation of select model genes, the abundance and distribution of such structures in the human genome has remained unclear. Here we utilize high affinity i-motif immunoprecipitation followed by sequencing to map i-motifs in human genomic DNA. Validated by biolayer interferometry and circular dichroism spectroscopy, our approach identified over 650,000 i-motif structures in human genomic DNA. The i-motif structures are widely distributed throughout the human genome and are common among highly expressed genes and in genes upregulated in G0/G1 cell cycle phase. Our findings provide experimental evidence for the widespread formation of i-motif structures in human genomic DNA.

---

Unravelling the location of regulatory elements in the human genome is critical for understanding genome architecture and function. Unlike the canonical double stranded B-form DNA [1], i-motif (iM) DNA is formed by hemi-protonated intercalated cytosine base pairs folded into a tetrameric structure. iMs and related guanine-rich G-quadruplex structures (G4s) have been identified as important regulatory elements in gene transcription, DNA replication, telomeric and centromeric regions and have been implicated in a range of human conditions [2-8]. While there has been extensive research into G4 location and function [9-13], detailed genomic insights into iM formation in human genomic DNA remains elusive.

## Immunoprecipitation and sequencing

To establish a map of iMs across the human genome, we first isolated genomic DNA from the MCF7 human breast cancer cell line previously utilized for G4 research [11, 14], and generated fragments of 100-200 bp through DNA shearing. Following a heat-cool step, immunoprecipitation of iM DNA was carried out using the previously developed high affinity iMab antibody [15] under physiological conditions. Six experiments were carried out (3 biological x 2 technical replicates) to generate annotations of iMs across the human genome. In total, we identified 683,100 i-motif regions (iMs) across three highly reproducible biological replicates (R^2^ > 0.89) (Fig. 1a, Extended data Fig. 1a). Sequences were analysed using MEME [16], which revealed the enrichment of cytosine rich motifs connected by thymidine tracks (Fig. 1b, Extended data Fig. 1b), in excellent agreement with previously described iM sequences [2, 17-19]. We observed widespread distribution of iMs throughout the human genome, including within intergenic regions, introns, exons and promoter regions (Fig. 1c).

**Fig. 1.**
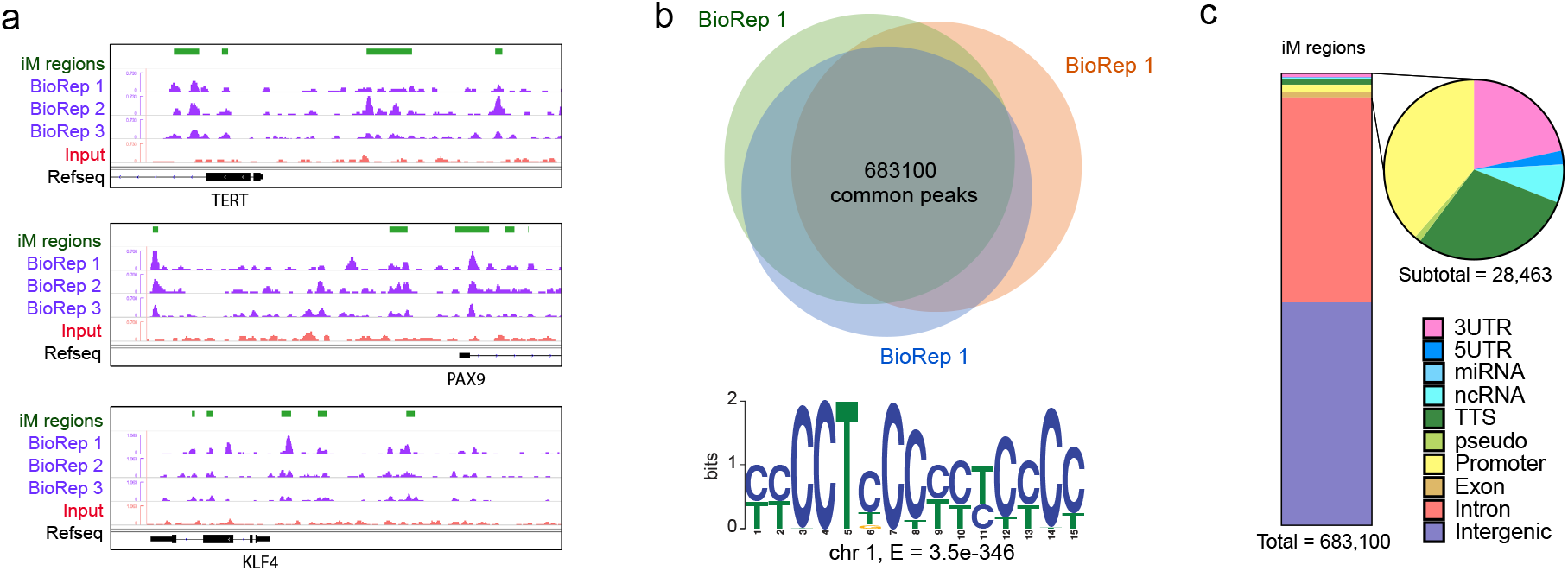
Distribution of iM structures across human genomic DNA. **a**, Genomic view highlighting iM structures in TERT, PAX9 and KLF4 oncogenes. iM regions from three biological replicates are shown (green, upper track: intersected iM regions; purple tracks: merged technical replicates for each biological experiment read values; lower red track: representative input profile). **b**, Total intersected iM peaks (683,103) of three biological replicates; most frequently identified sequence motif highlighted (chromosome 1 (MEME [16]). **c**, Distribution of iM structures across human genomic DNA.

## Biophysical validation

To further validate iM genomic sequences from identified regions and their ability to form iM structures, we selected 79 sequences located at the promoter regions of oncogenes and tumor suppressor genes (Fig. 2; Extended Table 1). We first confirmed the binding of the sequences to the iMab antibody by biolayer interferometry (BLI): this confirmed high affinity binding of 76/79 sequences with equilibrium binding constants (KD) ranging from 4 nM to 97 nM (Extended Table 1). Next, iM formation of a subset of the sequences was further validated by circular dichroism spectroscopy with the majority of analysed sequences displaying spectra indicative of iM structures (characterized by ∼285 nm /∼260 nm maximum/minimum [20], while control sequences did not display detectable iM spectra (Extended data Fig. 2a). For three of the iM structures and an hTelo positive control, folding was further investigated across a pH range of 5-8, with all of the analysed sequences displaying pH sensitivity, a canonical feature of the iM structure (Fig. 2, Extended data Fig. 2b) [21, 22].

**Fig. 2.**
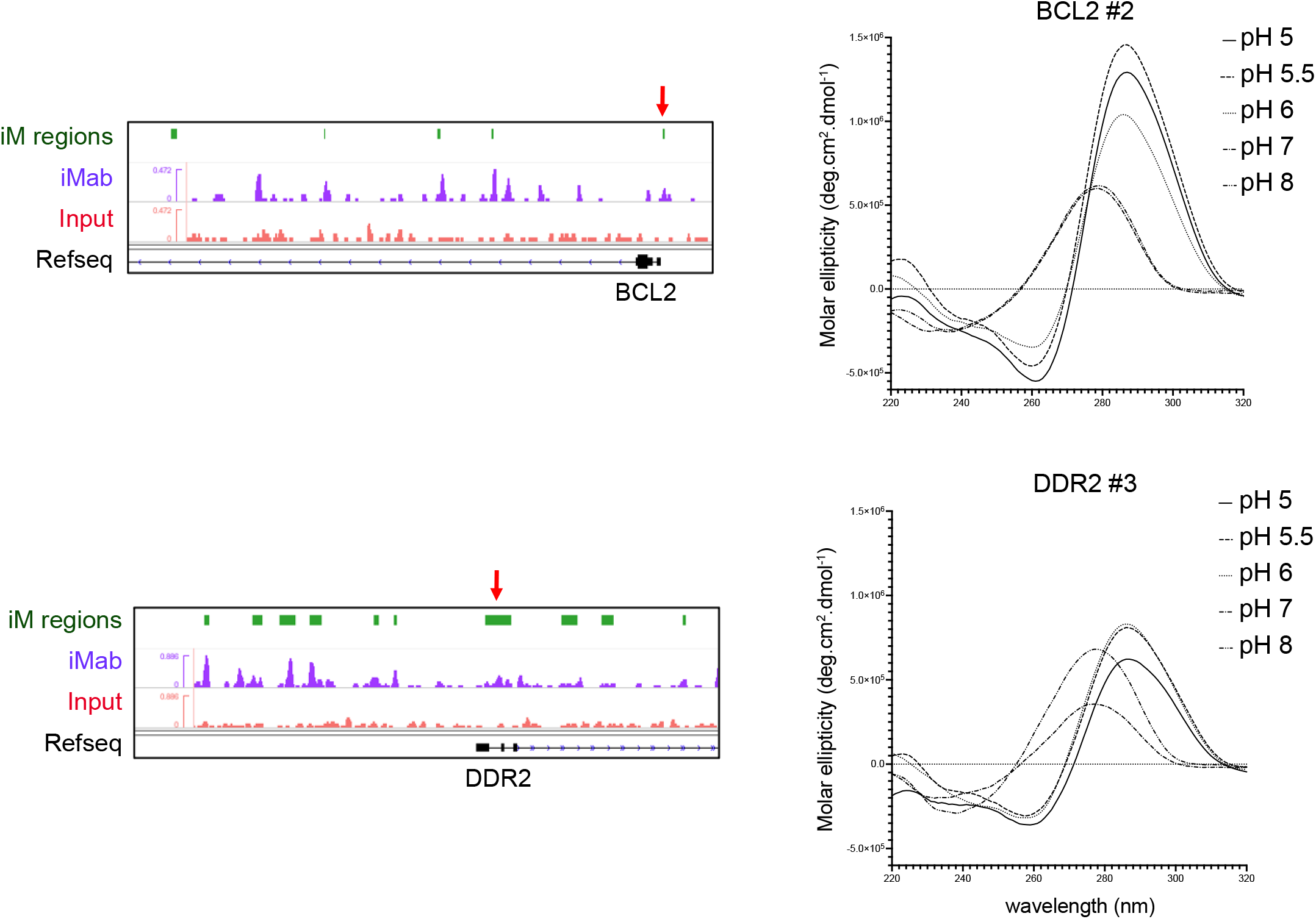
Evaluation of iM regions and biophysical validation. Genomic view highlighting identified iM structures in BCL2 and DDR2 genes. Representative iM regions from three intersected biological replicates are shown (upper green track: iM regions; purple track: representative merged biological replicate read values; lower red track: representative input profile). Validation of identified iMs upstream of the TSS (red arrow) by DNA synthesis and circular dichroism spectroscopy under variable pH conditions (pH 5 – 8; right panels).

## Interplay of G4 and iMs

Biochemical experiments have suggested a close relationship between G4 and iM formation [23]. It is also known that dynamic formation of either structure has direct effects on the physical properties of the opposing DNA strand and can lead to changes of genomic expression within neighbouring genes [24]. Using iMab we have recently demonstrated that G4 and iM formation are interdependent and that the stabilization of one structure can prevent the formation of the other [25]. Locations of G4 quadruplexes in human genomic DNA have been previously reported (using either PDS or K^+^ as stabilizing agents [9]), allowing for comparisons with the iM dataset generated here. An initial analysis revealed considerable colocalization between iM and G4 counts (Fig. 3a), indicating broad clustering of the two structures throughout DNA. Indeed, further analyses revealed that 34.2% of G4s reported previously after PDS stabilization overlapped with iMs (227,459/665,429) (Fig. 3b). Significant levels of overlap were observed despite the fact that the previously obtained G4 dataset was generated using polymerase stop assays [9], rather than the immunoprecipitation technique utilized here.

**Fig. 3.**
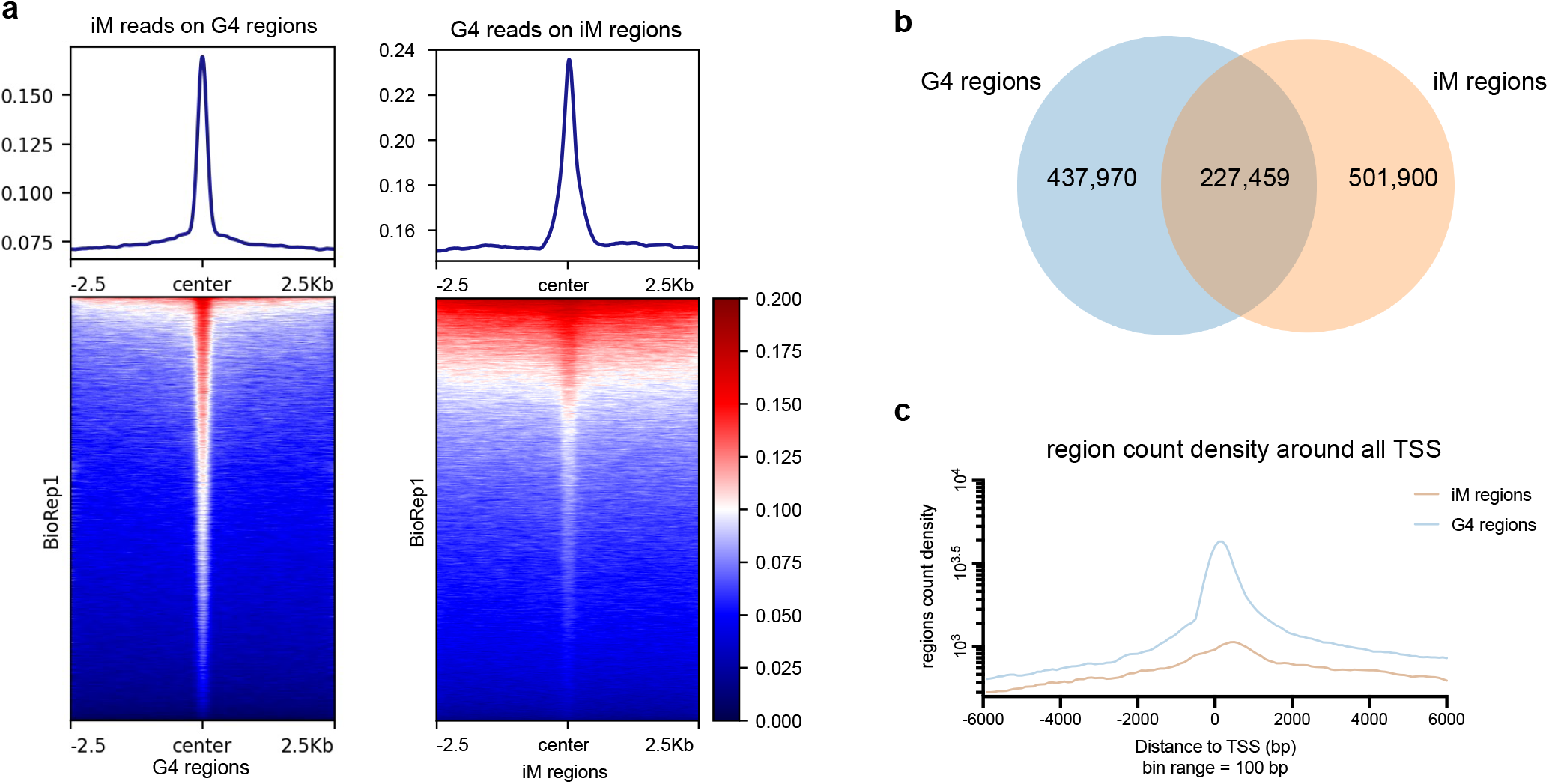
Comparison of iM and G4 annotations. **a**, Tag density histograms and heatmaps representing the occupancy of reads after iMab immunoprecipitation in proximity to published G4 regions [9] (left panel) and occupancy of G4 reads in proximity to iM regions (right panel) are shown Both datasets are the centred value of the site with 2.5Kbp flanks. **b**, Overlap of iM and G4 regions. **c**, Distance of iMs and G4s regions to proximal TSS (regions overlapping a TSS are noted as zero and count values given as the total density).

Previous studies have suggested wide ranging effect of iMs and G4s on transcription and gene regulation [12, 26, 27]. To further investigate this question, we determined their distance to the closest transcription starting site (TSS). For both DNA structures, we observed considerable overlap for all coding human genes (Fig. 3c), as indicated by very high site density counts in proximity to the TSS. This confirms biochemical studies of individual iMs [8, 24, 28-30] and computational predictions [31-33], that suggest close correlation between tetrameric DNA formation and transcriptional regulation. In addition to transcription, G4 structures have been implicated in the control of replication [3, 34]. When analysing the previously reported G4 dataset, we observed a clear association of G4s with early (but not late) replication domains. However, such association was not observed for the iM dataset, indicating differential roles of the two DNA motifs in replication (Extended data Fig. 3).

## iMs and gene expression

The accessibility of chromatin to transcription factors (TFs) and the regulation of gene expression is mediated by DNA accessibility and histone modification, including methylation and acetylation [35-37]. TF ChIP-seq data has been reported for multiple nuclear proteins in MCF7 cells [38-40], and we observed overlap of iMs with binding sites of transcription complex associated molecule including EP300, POLR2A, and CTCF (Extended data Fig. 4a). In addition to TFs, we observed overlap of iMs with histone modifications, including H3K4me1, H3K27ac, H3K4me3, as well as with open chromatin determined by DNase-seq and ATAC-seq. Although the variation in sample size of previously published datasets prevents more extensive analyses, positive correlation of IMs with H3K27me3 histone mark and several transcriptionally active regions was observed, exemplified by histone modification H3K27ac, and H3K4me3, while also negative correlation of iMs was observed for transcriptionally repressed regions, exemplified by H3K9me3 (Extended data Fig. 4b). High global read counts of TFs and histone modifications were further observed for iM regions, including POL2A, RAD51 and JUN. More specifically, within TSS and promoter regions, high read counts were observed in regions associated with active transcriptional regulation and open chromatin (Extended data Fig. 4c).

To further explore the association between iMs and gene expression, we surveyed previously published MCF7 RNA-seq datasets [41]. Our analyses revealed significant differences, with iMs most commonly observed among highly expressed genes (Fig. 4a), in agreement with the observed association of iMs with open chromatin and transcriptionally active DNA regions. Next, we investigated whether such iMs are associated with specific biological roles or gene families. Precisely focused our attention on a subset of genes containing high iM counts near their TSS region, indicating that these may be regulated by iM formation. Analyses from ontology associations particularly highlighted association with excitatory systems, including calcium and adrenergic signalling pathways, as well as genes involved in developmental pathways; significant associations were also observed with circadian entrainment pathways (Fig. 4b).

**Fig. 4.**
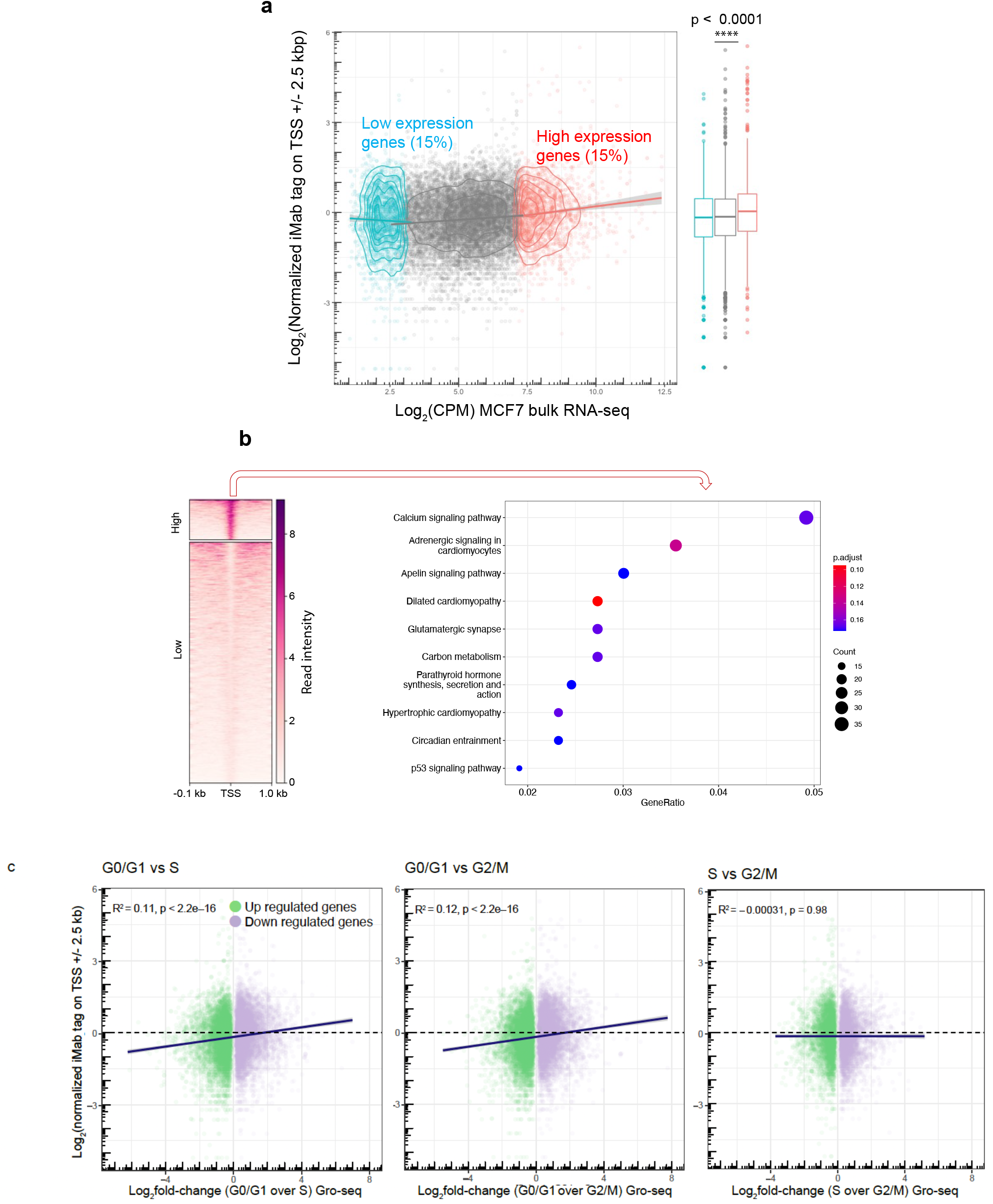
iMs are found preferentially around high expressed genes and genes upregulated during early cell growth phases. **a**, Scatter plot of Log_2_[counts per million] of expression values for published basal estate MCF7 cell and the Log_2_[normalized read tag on TSS +/-2.5Kbp]. The *x* axis indicates published data of basal estate (no hormone stimulated) MCF7 bulk RNA-seq counts, while the *y* axis indicates the normalized tag value around the same gene at a range of TSS +/-2.5Kbp. 15% of the upper and lower tier of gene expression values were assigned based on their Log_2_(CPM) as high expression or low expression. Replicated data for 14635 unique genes is shown, lines represent linear trends for each color group, low expression as blue, neutral expression as grey and high expression as red. Box plots indicate median values (centre lines) and IQRs (box edges), with whiskers ranging at lower quantiles -/+ (1.5 x IQR) and outliers beyond the range indicated as dots. (P < 0.0001 nonparametric One ANOVA Kruskal Wallis test). **b**, heatmap representing the results for the two subsets (high and low) from all refseq genes at the TSS region within a range of +/-1Kbp. Right panel shows the KEGG ontology analysis of genes with high iM read density across the TSS region. The 10 most significant molecular functions ortholog subsets are shown after a p < 0.5, *x* axis represents the gene ratio and the size of the dot show the total count of the group. **c**, Scatter plot of Log_2_[fold change] of expression values between G0/G1 and S, G0/G1 and G2/M or S and G2/M cell cycle phases in MCF7 cells and their relation against the Log_2_[normalized read tag on TSS +/-2.5Kbp] of iM reads. The *x* axis indicates reanalysed published data of MCF7 Gro-seq counts [43]. Purple dots indicate upregulated genes and green dots indicate downregulated genes (G0/G1 over S <2; n=5,360 upregulated and n=4,833 downregulated genes) (G0/G1 over G2/M < 2; n=4,775 upregulated and 4,689 downregulated) (S over G2/M <2; n=3,343 upregulated and n=3,530 downregulated genes). Non-significant genes were excluded from the plots.

Lastly, we analysed previously published nascent RNA (GRO-seq) based sequencing datasets from MCF7 cells [42, 43], focusing on differential RNA-seq of upregulated and downregulated genes within G0/G1, S, and G2/M cell cycle phases. These analyses revealed correlation of iM reads and upregulated genes in G0/G1 phase (G0/G1 vs S; R^2^=0.11; p < 2.2e-16, G0/G1 vs G2/M; R^2^=0.12; p < 2.2e-16, G2/M vs S; R^2^=-0.00031; p < 0.98) (Fig. 4d), consistent with previously reported data by our group and others that have demonstrated an increase of iM formation in G0/G1 phase by immunofluorescence [15, 25].

## Conclusion

Our study represents the first analysis of iM structures in human genomic DNA using immunoprecipitation and high-throughput sequencing. Using an iM specific antibody (iMab), we were able to experimentally map their location in human genomic DNA. The total number of iMs (683,100, as determined by immunoprecipitation) highly similar in scale to what has previously been reported for G4s (665,429, as determined by polymerase stalling [9]).Our study highlights the exquisite specificity of the iMab antibody fragment, not only for immunofluorescence [15, 25], but also for immunoprecipitation-based sequencing on a genome-wide scale, allowing the identification of a large number of iM forming sequences and their validation by biophysical characterization in this study. The use of the iMab antibody, developed in our laboratory, for iM immunoprecipitation is further supported by a recent study by Ma et al. [44] on transposable elements in rice; although the use of plant DNA and acidic conditions (pH 5.5) prevent direct comparisons, the study highlights the potential of iMab immunoprecipitation across a wide range of organisms.

We conclude that iM structures are located in close proximity to G4 forming regions, highly expressed genes and genes expressed in G0/G1 phase, highlighting their non-random distribution and involvement in genomic architecture. Our study provides foundational knowledge relating to the location and distribution of iMs in human genomic DNA, representing potential targets for future diagnostic and therapeutic strategies.

## Methods

### Antibody production

Expression and purification of the iM specific antibody (iMab) was performed as previously described [15, 45]. In brief, the iMab gene was cloned into a pET12a vector encoding C-terminal c-Myc and Avi tags. Competent *E. coli* BL21-Gold (DE3) cells were transformed with pET12a-iMab and pBirAcm (Avidity). *In vivo* biotinylated iMab-MycAviTag antibody fragments were purified by affinity chromatography and biotinylation confirmed by biolayer interferometry using a BLItz system (Pall ForteBio LLC) and streptavidin sensors.

### DNA immunoprecipitation and Sequencing

Genomic DNA from MCF7 cells was harvested using an AllPrep DNA/RNA/miRNA Universal Kit (QIAGEN, Cat No: 80224) following the manufacturer protocol. Purified DNA was concentrated using Amicon Ultra-0.5 Centrifugal Filter Unit (Millipore, UFC501096) and concentration adjusted. DNA was fragmented using AFA Fiber Pre-Slit Snap-Cap microtubes (Covaris, 520045) and a Covaris instrument (M220 Focused-ultrasonicator) at Peak power: 50, Duty factor: 20, Cycles/Burst: 200, Time: 700 seconds, Setpoint: 20 °C producing 100 – 200 bp fragments. DNA fragments were stored at - 80 °C until a pull-down experiment. A 96-well MaxiSorp plate (Thermo Scientific, 442404) was coated by adding 60 *µ*L per well of 50 *µ*g mL^-1^ streptavidin diluted in PBS, and incubated overnight at 4 °C. Fragmented DNA was thawed and diluted in DPBS (ThermoFisher, 14190144), pH adjusted to 7.4. For each biological replicate, 140 *µ*g DNA were diluted in 525 *µ*L DPBS and heated at 90 °C for 10 minutes followed by cooling down at the rate of 1 °C per minute to 21 °C and kept on ice until the pull-down experiment. A streptavidin coated plate was washed once with PBS and blocked with 200 *µ*L per well of SuperBlock (ThermoFisher, 37515) for two hours at room temperature. Biotinylated iMab-MycAviTag was diluted to 49 *µ*g in 525 *µ*L of DPBS pH 7.4 (7 x wells, 75 *µ*L each) for each biological replicate. Blocked wells were washed once with PBS and Biotinylated iMab-MycAviTag solution was added. After incubating for 1 hour at room temperature while shaking at 200 rpm, wells were washed twice with DPBS pH 7.4. Then, 75 *µ*L of DNA solution was added to each well and incubated 16 – 20 hours at 4 °C. The supernatant was aspirated, each well washed and tap-dried seven times with 200 *µ*L of DPBS pH 7.4 supplemented with 0.1 % Tween 20 and twice with DPBS only. Elution was performed by adding 100 *µ*L of 100 mM Tris/Acetate pH 10 supplemented with 1 % SDS to each well and incubating for one hour at 40 °C while the plate was shaking at 200 rpm.

Eluted DNA were collected, combined, and concentrated using ssDNA/RNA Clean & Concentrator kit (Zymo, Cat No: D7011) following the manufacturer protocol. Accel-NGS 1S Plus DNA kit (Swift Bioscience, Cat No: 10024) and 1S Plus Dual Indexing kit (Swift Bioscience, Cat No: 18096) were used to prepare a library compatible with illumina sequencing platform. A 4200 TapeStation instrument (Agilent Technologies) and D1000 ScreenTapes (Agilent Technologies, Part No: 5067-5582) with D1000 Reagents (Agilent Technologies, Part No: 5067-5583) were used for library quality controls. Sequencing was performed on NextSeq 500 instrument (illumina) using NextSeq 500/550 Mid Output Kit v2 (illumine, FC-404-2001) in a paired-end 75 x cycles mode following the manufacturer protocol. At least 25 % spike-in PhiX control V3 (illumine, 15017666) was added to the final solution prepared for sequencing.

### iM analyses

Fastq files were trimmed by removing 5 bp and 10 bp from the 5’ end of forward and reverse reads, respectively, using cutadapt (v 1.16) [46]. Quality control was performed on the trimmed reads using FastQC software (https://www.bioinformatics.babraham.ac.uk/projects/fastqc/). ENCODE Uniform Processing pipeline (paired-end, replicated TF ChIP-seq, hg19) available on DNAnexus was used to call broad peaks. Each biological replicate comprises of two technical replicates. The same input DNA (two technical replicates) was used for the peak calling of biological replicates. Broad peaks from three biological replicates were interested using BEDTools (v 2.27.0) [47] and a subset of peaks with length between 25 to 5000 bp were used for the downstream analyses. Peaks were annotated using HOMER (v 4.11.1) [48]. Regions were transformed back to fasta using bedtools -getfasta from reference genome hg19, sequences under 30bp discarded using seqkit and de novo motif discovery was performed using MEME (v 5.3.3) [16] while 0-order model of sequences was used as the background model with the length between 11 to 49 bp.

### DNA synthesis and biolayer interferometry

iM DNA sequences were generated by DNA synthesis (Integrated DNA Technologies) and resuspended at 100 µM in nuclease-free mQ-H_2_O. Binding kinetics of iMab and iM DNA oligonucleotides were determined by biolayer interferometry using a BLItz system (ForteBio) at room temperature. Biotinylated iMab scFv (or DP47/DPK9 as control) in 20 mM Tris-Acetate pH 6.0 + 100 mM KCl was used to load the streptavidin biosensor for 2 min. Refolded DNA oligonucleotides in 20 mM Tris-Acetate pH 6.0 + 100 mM KCl at 100 nM were then used for association for 4 min, followed by 4 min dissociation in 20 mM Tris-Acetate pH 6.0 + 100 mM KCl. Curve fitting was performed using BLItz Pro software version 1.3.

### Circular Dichroism Spectroscopy (CD)

For characterization, DNA oligonucleotides were diluted to 10 μM in 20 mM Tris-Acetate pH 6.0 + 100 mM KCl and annealed by heating at 90 °C for 10 min and cooling down to room temperature at the rate of 1 °C per min. CD spectra were recorded using an Aviv 215S circular dichroism spectrometer equipped with a Peltier temperature controller at 25 °C. Four scans were gathered over the wavelength ranging from 220 to 320 nm in a 0.1 cm path length cell at the standard sensitivity, data pitch 0.1 nm, continuous scanning mode, scanning speed 100 nm min^-1^, response 4 s, and bandwidth 1 nm. 20 mM Tris-Acetate pH 6.0 + 100 mM KCl was scanned as buffer sample and its spectrum subtracted from the average scans for each sample. CD spectra were collected in millidegrees, normalized to the total species concentrations, and stated as molar ellipticity units (deg.cm^2^.dmol^-1^). Each CD spectrum was smoothed by averaging ten neighbour points using GraphPad Prism software. For pH gradient folding, DNA oligonucleotides were diluted to 10 μM in 20 mM Na-cacodylate at pH 5.0, pH 5.5, pH 6.0, pH 7.0 or pH 8.0 + 100 mM KCl and annealed by heating at 90 °C for 10 min and cooling down to room temperature at the rate of 1 °C per min, as described [20]. CD spectroscopy was performed as previously described, using the specific pH buffer scans for subtraction.

### Analyses of G4-seq and ChIP-seq data

G4-seq data and multiple datasets from ChIP sequencing, ATAC and DNase seq from MCF7 cells were obtained from previously published data and analysis performed against iMs using bedtools [47], and deepTools [49] version 3.5.0 to transform data when necessary and plot. Analyses used regions within a ± 1kbp from the TSS of all refseq [50] genes, defined as TSS proximal iM regions. Read heatmaps and correlation plots were drawn using deepTools [49] and R version 4.0.4. Ontology analyses were done in R using the packages: ChIPseeker [51], TxDb.Hsapiens.UCSC.hg19.knownGene, EnsDb.Hsapiens.v75, clusterProfiler, AnnotationDbi, and org.Hs.eg.db,

### Analyses of RNA-seq and GRO-seq data

RNA-seq and GRO-seq read counts were obtained from previously published data on unstimulated MCF7 cells [41, 43]. Reads were annotated, depleted for low expressing genes (CPM < 0.5), normalized for composition bias and merged with iMab tag on TSS +/-2.5 kb or tested for differential expression analysis. All reanalysis was conducted in R using the packages: EdgeR, limma, Glimma and org.Hs.eg.db.

### Statistical Analyses

Statistical tests were collected using deepTools [49], R version 4.0.4 and Prism 9. The ranges of *x* and *y* axes for scatter plots are justified in the figure captions. Statistical tests, number of experiments and *P* values are cited in figures and/or captions.

## Data availability

Sequencing data used in this manuscript can be accessed through the Gene Expression Omnibus under the accession code (XXXXXXXX). Specially, G4-seq data was used from previously reported data with accession code GSE63874. MCF7 public referenced data for analyses are available from accession codes GSE47042 (RNA-seq), GSE94479 (GRO-seq), GSE53984 (Origins of replication), GSM816627 (DNase-seq), GSM4130902 (ATAC-seq), ChIP sequencing data from multiple proteins was obtained from GSM822295 (POLR2A), GSE96352 (H3K27ac), ENCFF783ZIE (H3K4me3), GSE86714 (H3K4me1), ENCFF000VGA (H3K27me3), ENCFF350GNM (H3K9me3), GSM822305 (CTCF), GSE91550 (JUN), GSM1010892 (JUND), GSE105597 (RAD51), GSE55921 (BRD4), GSM1010800 (EP300), and CpG islands (UCSC hg19 genome annotation).

**Extended Figure 1.**
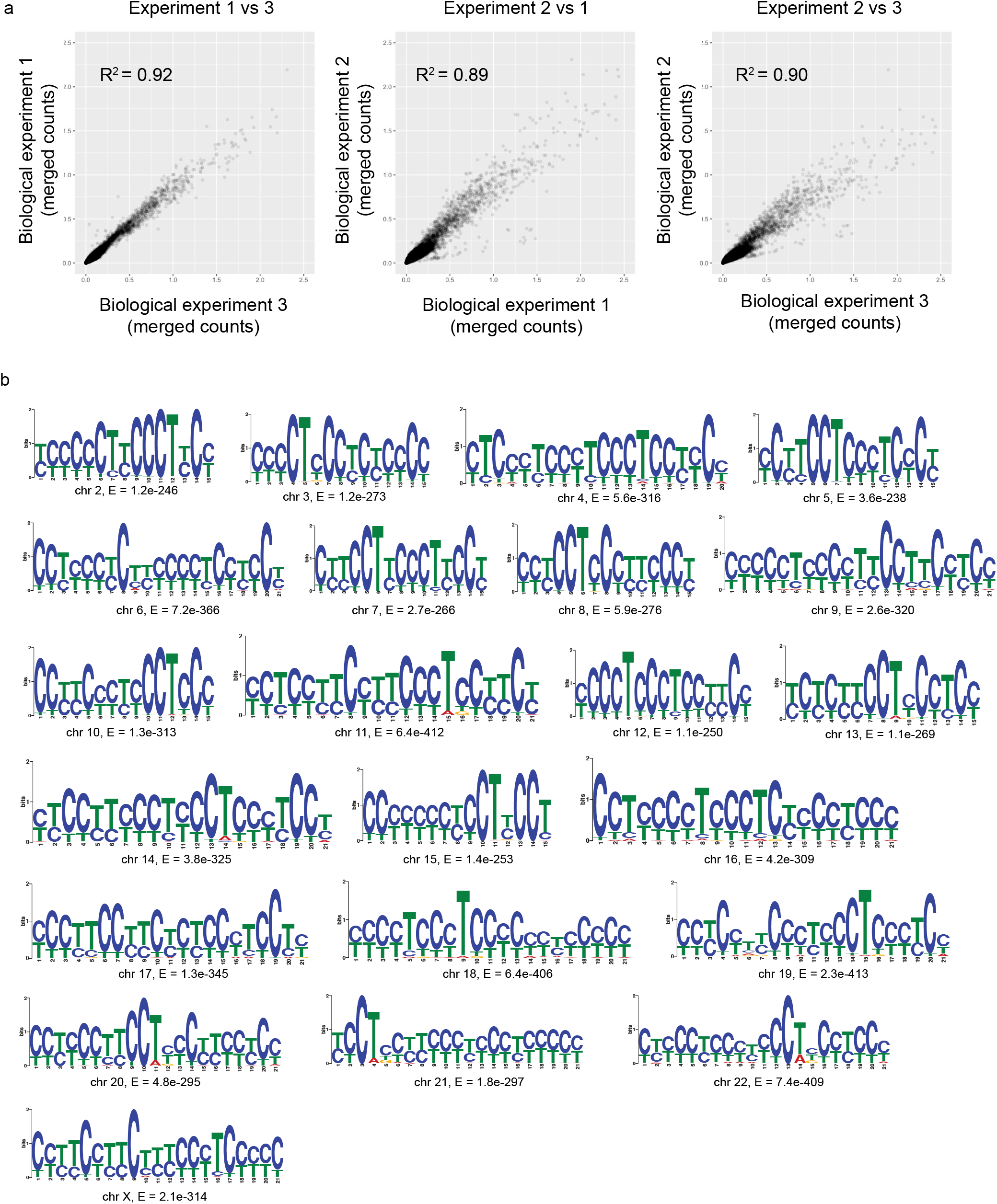
iM immunoprecipitation and commonly observed iMs. **a**, Pairwise comparison of tag counts after iMab immunoprecipitation for each biological replicate (Pearson correlation between two experiments shown; deepTool; bamCompare using default parameters). **b**, Most frequently identified iM sequence motifs (MEME [16]).

**Extended Table 1.**
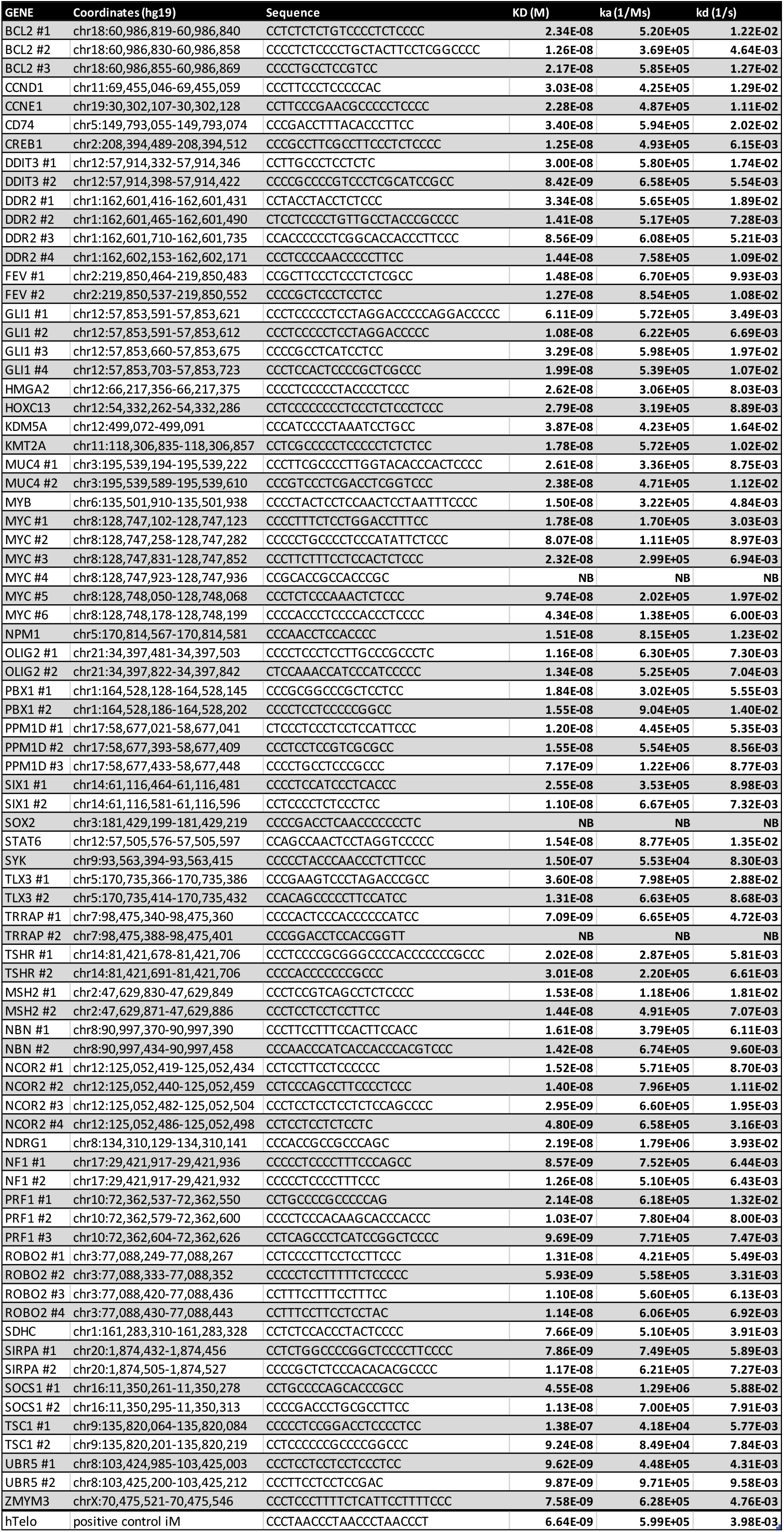
Oligonucleotides used for biolayer interferometry (BLI) and circular dichroism (CD) studies. Gene name, genomic position and affinity (as determined by BLI) with KD, ka and kd reported. NB: non-binding.

**Extended Figure 2.**
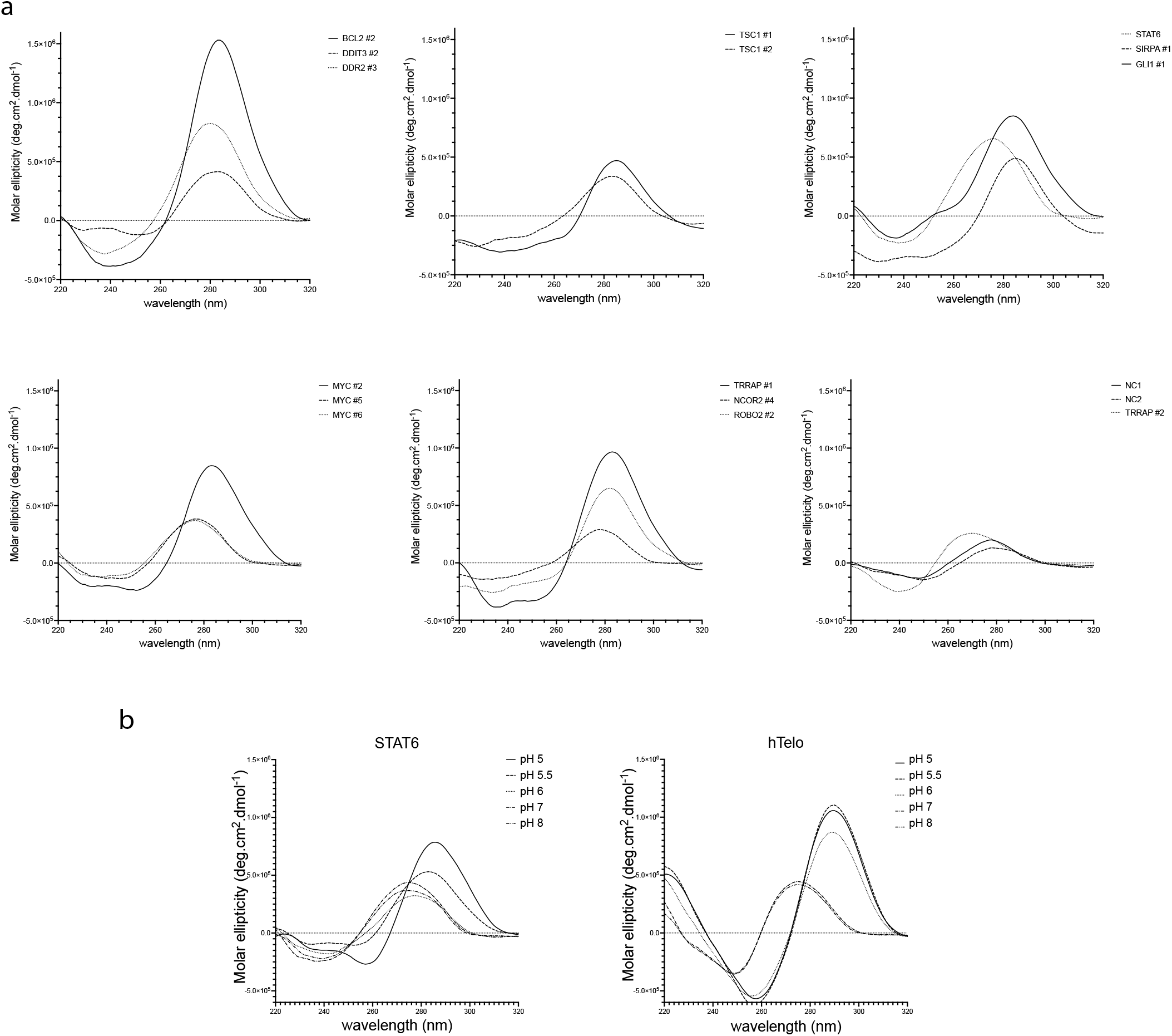
Biophysical validation. **a**. Validation of identified iMs in BCL2, DDIT3, DDR2, TSC1, STAT6, SIRPA, GLI1, MYC, TRRAP, NCOR2 and ROBO2 by DNA synthesis and circular dichroism spectroscopy at pH 6.0. TRRAP #2 did not display detectable iMab binding (Extended Table 1) and was used as a negative control, in addition to NC1(5-‘CAGACTGTCGATGAAGCCCTG-3’) and NC2 (5’-CTAGTTATTGCTCAGCGGTG-3’) negative control sequences. b. Validation of identified iM in STAT6 by DNA synthesis and circular dichroism spectroscopy at pH 5-pH8 range with hTelo [22] shown as a positive control.

**Extended Figure 3.**
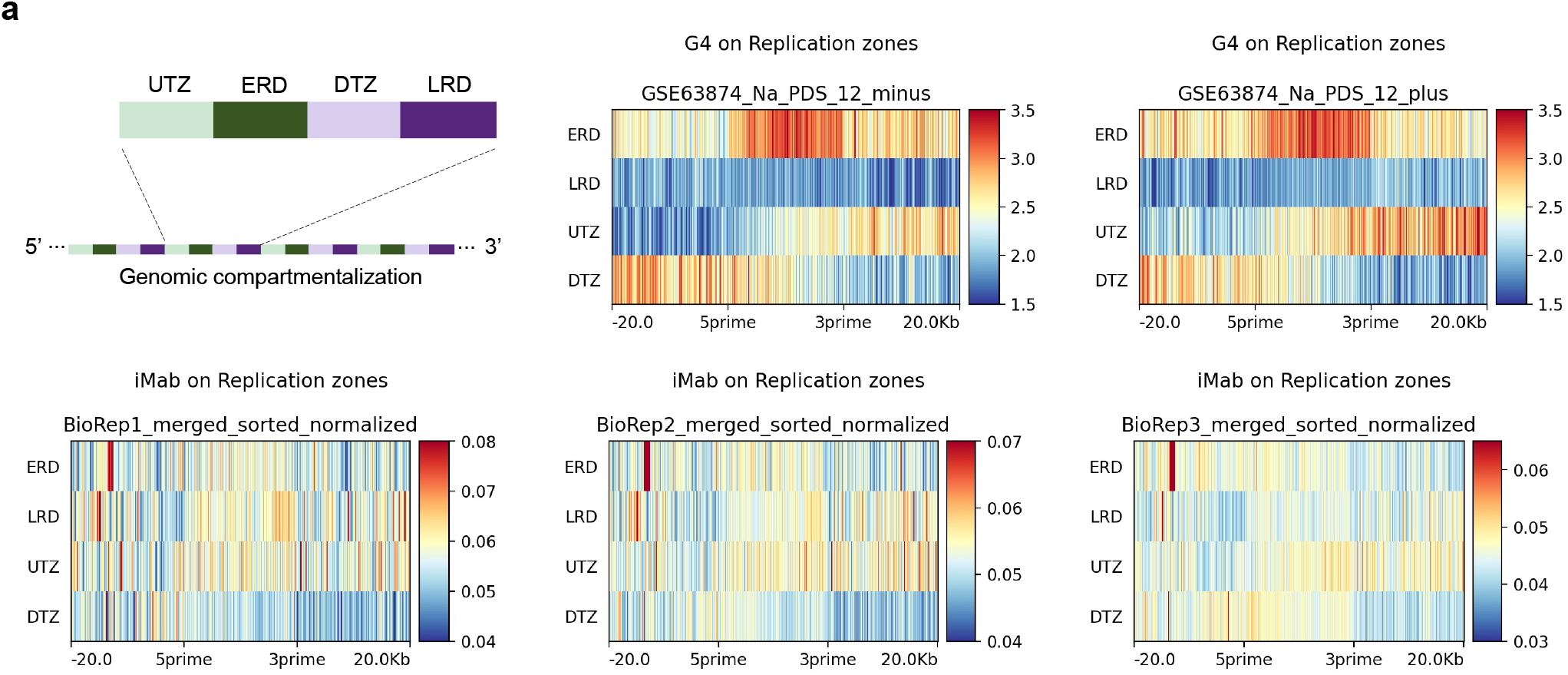
Comparison of iM and G4 annotations within replication zones. Genomic partition of replication zones (analyses based on repetition of up transition zones (UTZ), early replication zones (ERD), down transition zones (DTZ) and late replication domains (LRD); Tag count heatmaps of G4 [9] (upper panels) and iMs (lower panels; three biological replicates shown).

**Extended Figure 4.**
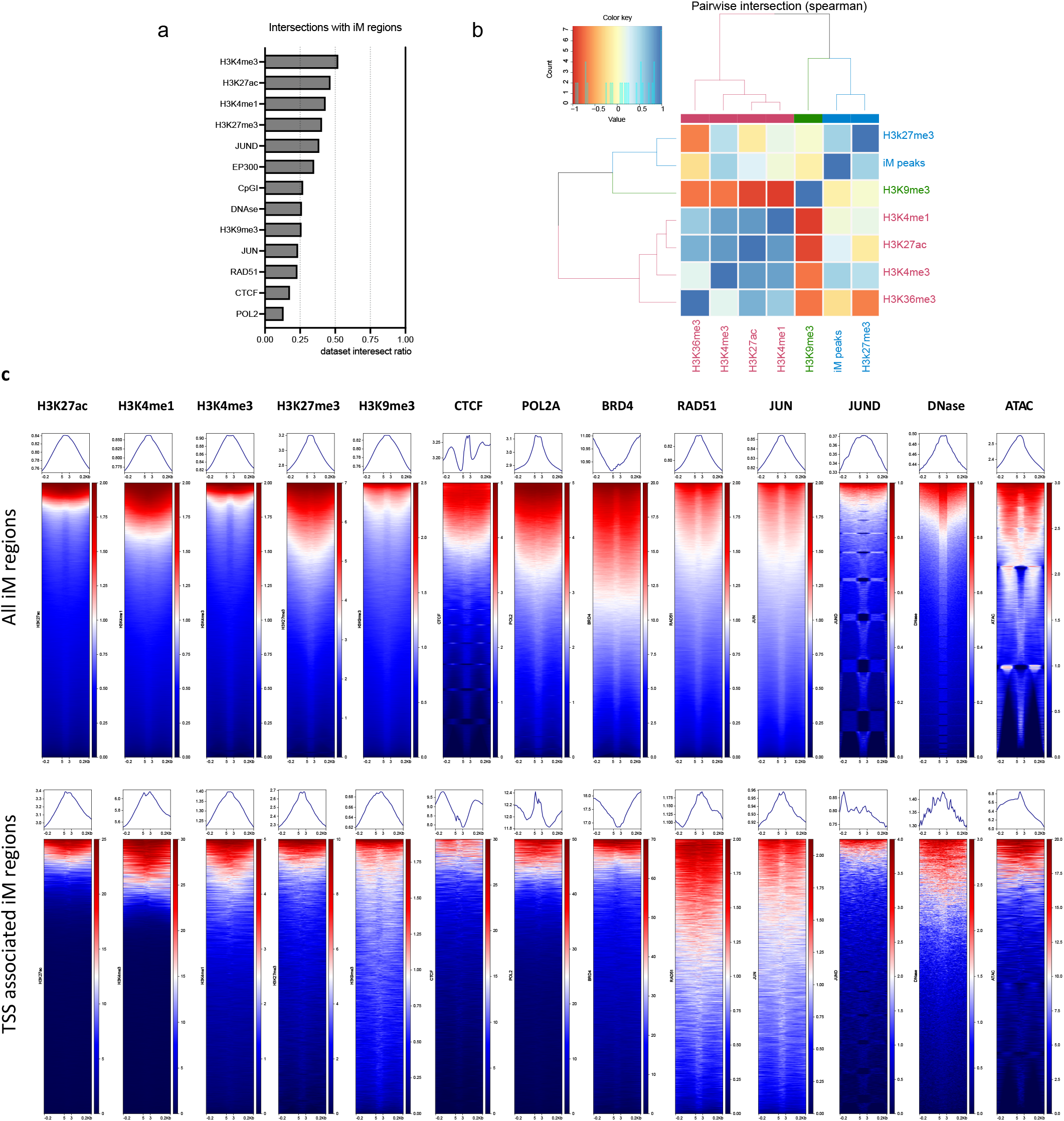
iM regions occupy transcription active sites. **a**, Intersection of transcription factor binding sites and histone marks with iM regions (see Methods for detail). **b**, Pairwise intersection of iM regions and histone marks (spearman correlation values shown). **c**, Common histone modification marks and transcription factors in iM regions (top panels: all iM regions; bottom panels: TSS-promoter associated regions).

